# Pili Expression in *Geobacter sulfurreducens* Lacking the Putative Gene for the PilB Pilus Assembly Motor

**DOI:** 10.1101/2021.07.10.451916

**Authors:** Toshiyuki Ueki, David J.F. Walker, Kelly P. Nevin, Joy E. Ward, Trevor L. Woodard, Stephen S. Nonnenmann, Derek R. Lovley

## Abstract

Multiple lines of evidence suggest that electrically conductive pili (e-pili) are an important conduit for long-range electron transport in *Geobacter sulfurreducens*, a common model microbe for the study of extracellular electron transport mechanisms. One strategy to study the function of e-pili has been to delete the gene for PilB, the pilus assembly motor protein, in order to prevent e-pili expression. However, we found that e-pili are still expressed after the gene for PilB is deleted. Conducting probe atomic force microscopy revealed filaments with the same diameter and similar current-voltage response as e-pili harvested from wild-type *G. sulfurreducens* or when e-pili are heterologously expressed from the *G. sulfurreducens* pilin gene in *E. coli*. Immunogold labeling demonstrated that a *G. sulfurreducens* strain expressing e-pili with a His-tag continued to express His-tag labelled e-pili when the PilB gene was deleted. Strains with the PilB gene deleted produced maximum current densities comparable to wild-type controls. These results demonstrate that deleting the gene for PilB is not an appropriate strategy for constructing strains of *G. sulfurreducens* without e-pili, necessitating a reinterpretation of the results of previous studies that have employed this approach.

**Importance:** *Geobacter sulfurreducens* is a model microbe for the study of biogeochemically and technologically significant processes such as the reduction of Fe(III) oxides in soils and sediments; bioelectrochemical applications that produce electric current from waste organic matter or drive useful processes with the consumption of renewable electricity; direct interspecies electron transfer in anaerobic digestors and methanogenic soils and sediments; and metal corrosion. The phenotypes associated with gene deletions are an important strategy for determining the mechanisms for extracellular electron transfer in *G. sulfurreducens*. The results reported here demonstrate that a gene deletion previously thought to prevent the expression of electrically conductive pili in *G. sulfurreducens* does not have the intended result. Conductive pili continue to be expressed. This finding is important for interpreting the results of several previous studies on conductive pili function.

## Introduction

Strains of *Geobacter sulfurreducens* and closely related *Geobacter* species produce some of the highest recorded current densities in microbial fuel cells and *Geobacter* species are often the most abundant microorganisms within anode biofilms harvesting current in open systems such as sediments or wastewater (1, 2). *G. sulfurreducens* has also served as a convenient model microbe for other extracellular electron transfer process such as Fe(III) oxide reduction, direct interspecies electron transfer, and corrosion (1, 3, 4). However, developing definitive models for extracellular electron transfer in *G. sulfurreducens* has been challenging. *G. sulfurreducens* produces a wide diversity of outer-surface redox-active proteins and electrically conductive filaments and can rapidly adapt expression of those outer-surface components in response to selective pressure that favors specific types of extracellular electron transfer, or to mutations that disable function of one or more of the outer-surface components (3, 5–8). Gene deletions designed to elucidate function may have unintended negative impact on the expression of non-target proteins (1).

For example, multiple lines of evidence suggest that electrically conductive pili (e-pili) are essential for long-range electron transport through current-producing biofilms, (9, 10), but e-pili function cannot be reliably determined simply by deleting the gene for PilA, the monomer pilin protein that assembles into e-pili because the expression of other outer-surface proteins, such as multi-heme *c*-type cytochromes can be negatively impacted (11, 12). Furthermore, *G. sulfurreducens* pili may have additional functions, such as aiding in attachment to surfaces and biofilm formation (13). One strategy for avoiding these complications is to replace the gene for PilA with a gene that encodes a pilin that assembles into a poorly conductive pilin (14, 15). This strain, designated Aro-5, produces maximum currents much lower than wild-type cells and comparable to the *pilA* deletion mutant (14). In contrast to the *pilA* deletion mutant, which forms thin biofilms on current-harvesting electrodes (16), strain Aro-5 produces thick biofilms, but with much lower conductivity than wild-type biofilms (14).

An alternative strategy, proposed to avoid concerns of proper outer-surface cytochrome expression, is to delete the gene encoding PilB, the putative pilus assembly motor protein (12). However, the proposed lack of e-pili expression was only inferred from the expected impact of a *pilB* deletion on pili expression, not directly demonstrated (12). The PilB-deficient strain produced a maximum current (0.52 mA) that was half that of the wild-type strain (1.06 mA). This is a substantially higher proportion of wild-type current production than achieved by the Aro-5 strain, which only generated maximum currents ca. 15% of wild-type (14). The current production phenotype of the *pilB* deletion was evaluated in ‘batch mode’ in which the supply of electron-donor substrate is provided only at the beginning of the incubation and is depleted over time. It is well known that maximum current production in *G. sulfurreducens* is best evaluated when cells are grown in anode chambers in which fresh medium is continuously supplied once current production is (16). This method avoids the possibility of current production being prematurely inhibited due to a lack of electron donor or other medium constituents, or an accumulation of toxic byproducts over time (17).

In fact, a subsequent study (18) presented data that suggested that deleting the same PilB gene did not meaningfully affect current production. The *pilB*-deficient strain had an initial lag period that was ca. 10 % longer than wild-type (ca. 88 h versus 80 h) prior to the initiation of rapid current increase (Figure 1a). However, lag times are inherently variable even within replicates of the wild-type strain and slight differences in the initial start of current production are not meaningful phenotypes for interpreting long-range electron transport mechanisms, which are only required after thick biofilms begin to form (19). Unfortunately, the incubation reported (18) was not long enough to determine whether the PilB-deficient strain produced the same maximum current as the wild-type strain. These results suggested that further evaluation of the impact of deleting *pilB* was warranted.

**Figure 1.**
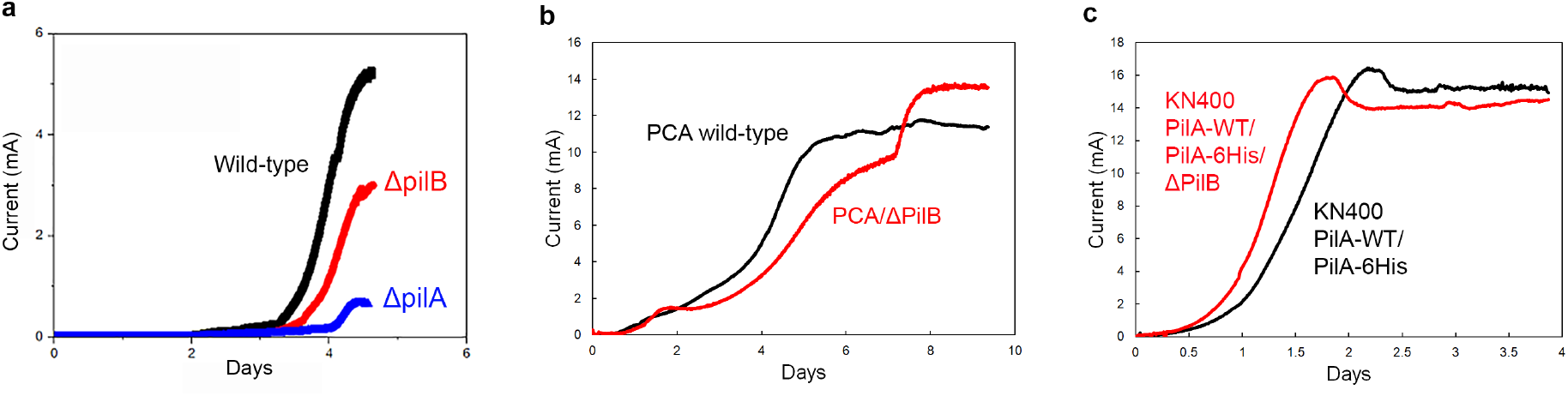
Current production *G. sulfurreducens* strain PCA. (a) Data image from (18) modified only to label each of the different colored curves. (b) Current production for *G. sulfurreducens* strain PCA wild-type and *G. sulfurreducens* strain PCA/ΔPilB. (c) Current production for *G. sulfurreducens* KN400 strain PilA-WT/PilA-6His and KN400 strain PilA-WT/PilA-6His/ΔPilB. Image in panel a is reproduced with permission.

## Results and Discussion

### High current densities and e-pili expression in Δ*pilB* in strain PCA background

In order to further evaluate the impact of deleting *pilB*, the same gene previously studied (12, 18) was deleted from *G. sulfurrreducens* strain PCA. This strain was designated ΔPilB. Current production was evaluated in anode chambers in which fresh medium was continuously supplied once current production initiated as previously described (17). Under these optimized conditions, strain ΔPilB produced current comparable to the wild-type (Figure 1b).

Multiple studies have demonstrated that *G. sulfurreducens* only produces high current densities when it is expressing e-pili (9, 14, 20, 21). Heterologous expression of pilin genes that yield poorly conductive pili consistently result in lower current production. Thus, the high current densities in strain ΔPilB were consistent with e-pili expression. Furthermore, transmission electron micrographs of strain ΔPilB revealed filaments that appeared to be pili emanating from the cells (Figure 2a).

**Figure 2.**
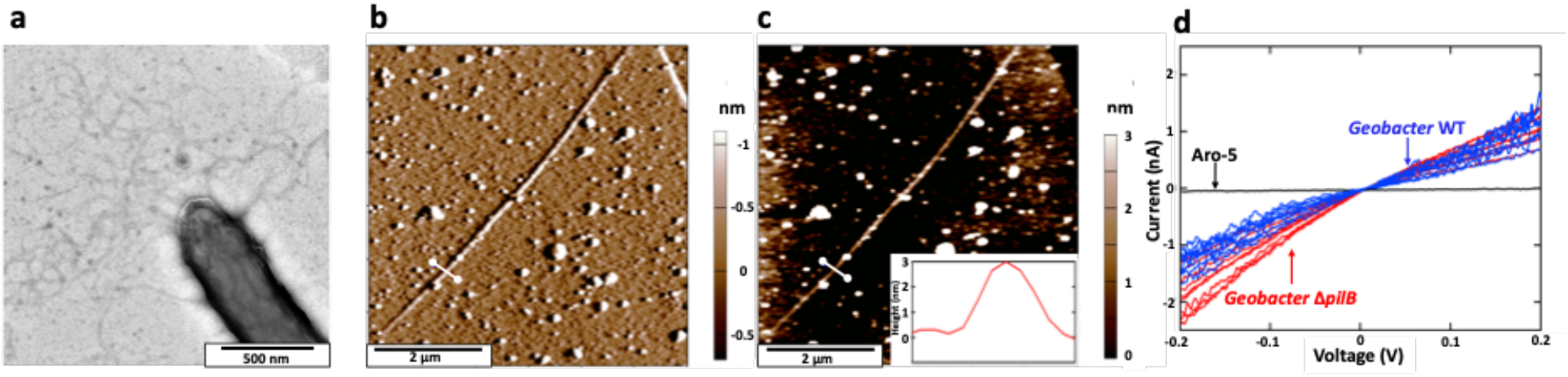
Expression of electrically conductive pili in the strain ΔPilB. (a) Transmission electron micrograph of negatively stained cell. (b) Atomic force microscopy, non-contact topographical imaging with deflection output of a single pilus, harvested from cells, laying on highly oriented pyrolytic graphite. (c) Height of the filament at the white line cross-section shown on the filament with the height profile shown in the inset. (d) Point-mode current response (I-V) spectroscopy of the individual *G. sulfurreducens* strain ΔPilB filaments shown in red overlaid with wild-type *G. sulfurreducens* in blue and *G. sulfurrreducens* strain Aro-5 in black. Data for pili from wild-type and Aro-5 strain are from reference (25).

Therefore, pili harvested from strain ΔPilB were examined with conducting probe atomic force microscopy, as previously described (22). Single filaments deposited on highly oriented pyrolytic graphite, which were identified in non-contact mode (Fig. 2b) had a height (Fig. 2c) of 3.0 ± 0.09 nm (mean ± standard deviation n=18; 6 individual points on 3 individual pili), consistent with the diameter of pili expressed in wild-type *G. sulfurrreducens* and the pili expressed when the *G. sulfurreducens* PilA pilin gene is heterologously expressed in *Pseudomonas aeruginosa* (23) or *Escherichia coli* (24). The response of individual filaments in point mode current response (I-V) spectroscopy was similar to that previously described for the e-pili of wild-type *G. sulfurrreducens* (Fig. 2d) and the e-pili produced when the *G. sulfurreducens* PilA is heterologously expressed in *E.coli* (24). The calculated conductance was 7.07 ± 0.81 nS (mean ± mean n=9; 3 individual points on 3 individual pili). These results suggested that strain ΔPilB generated high current densities because it expressed e-pili.

### Evaluation of *pilB* deletion in a strain expressing His-tagged pilin

In order to further evaluate the impact of deleting the PilB gene on e-pili expression in *G. sulfurreducens* we took advantage of the availability of a genetically modified strain of *G. sulfurreducens* KN400 designated PilA-WT/PilA-6His (26). Wild-type *G. sulfurreducens* KN400 offers advantages for the study of the role of e-pili in anode biofilms because: it produces higher current densities than strain PCA (27), with more pili (27) and higher biofilm conductivities (28, 29) than strain PCA (27). The PilA-WT/PilA-6His strain of KN400 expresses a wild-type PilA pilin monomer and a PilA monomer modified with a ‘His-tag’ (six histidines) at the carboxyl end (26). These pilin monomers assembled into e-pili emanating from cells are the e-pili are readily identified with an immunogold/TEM procedure that employs an antibody that recognizes the His-tag (26). The e-pili with the His-tag have conductivities comparable to wild-type e-pili and the strain expressing the His-tag e-pili produces current densities comparable to the wild-type strain (26).

The *G. sulfurreducens* KN400 gene encoding the same protein previously identified as PilB (12) in strain PCA of *G. sulfurreducens* was deleted from the KN400 strain PilA-WT/PilA-6His to yield strain KN400/PilA-WT/PilA-6His/ΔPilB. His-tag immunogold labeling of strain PilA-WT/PilA-6His/ΔPilB revealed labelled filaments (Figure 3) similar to those previously reported (26) for the strain without the *pilB* deletion. As noted previously for the strain without the *pilB* deletion (26), no unlabeled filaments were observed suggesting that e-pili were the primary filaments expressed. These results demonstrated that e-pili comprised of PilA monomers continued to be expressed when *pilB* was deleted. Further evidence for this was that strain PilA-WT/PilA-6His/ΔPilB produced current as well as strain PilA-WT/PilA-6His (Figure 1c).

**Figure 3.**
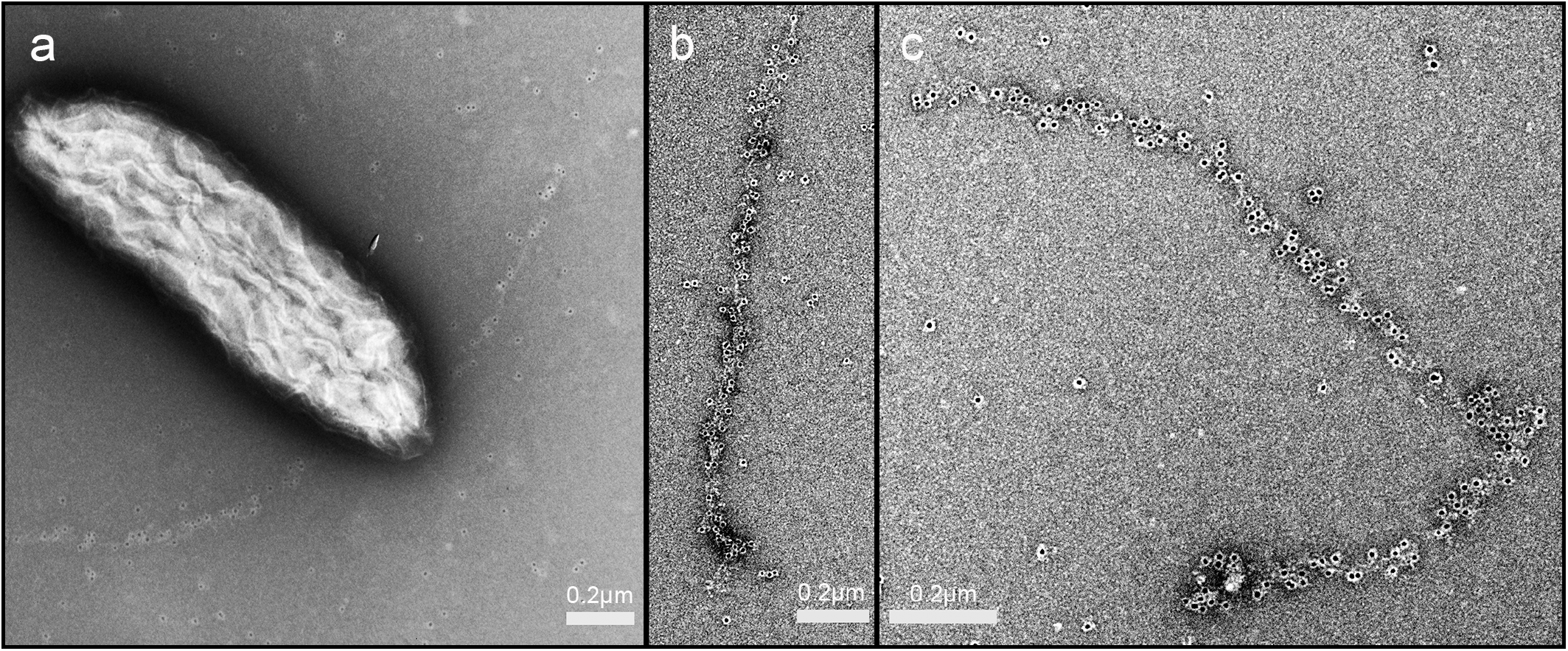
Immunogold labeling of pilin-containing filaments from strain PilA-WT/PilA-6His/ΔPilB. (a) Labeled filaments emanating from cell. (b and c) higher magnification of labelled filaments.

These results demonstrate that the assumption in previous studies (12, 18) that deleting the gene for the putative PilB in *G. sulfurreducens* prevents expression of its e-pili is not warranted. *G. sulfurreducens* PilB is homologous to other bacterial PilB ATPases and contains highly conserved features such as Walker A and B motifs, a Asp box, a His box, Arg fingers, and a tetracysteine Zn^2+^-binding motif (30). The gene (GSU1491) for PilB is located in a cluster including *pilT-4*, *pilC*, *pilS*, *pilR*, and *pilA* on the *G. sulfurreducens* genome (31). However, there are multiple genes within the *G. sulfurreducens* genome that might function as a PilB ATPase, or compensate for deletion of the gene for the PilB ATPase, which were previously reported (32) to be highly expressed in current-producing biofilms (Table 1). Another possibility is that type IV e-pili could be assembled by a type II protein secretion system, which is similar to type IV pilus assembly system in composition and structure and polymerizes pseudopilins (33, 34).

**Table 1.** PilB homologs in *G. sulfurreducens*.

| Gene | Description | Identity | Transcript abundance |
| --- | --- | --- | --- |
| GSU1491 | Type IV pilus biogenesis ATPase PilB | 100% | 9.91 |
| GSU0328 | Type II secretion system ATPase GspE | 45% | 10.12 |
| GSU0435 | PilB/PulE/GspE family ATPase | 26% | 6.86 |
| GSU1783 | Type II secretion system ATPase PulE | 38% | 9.70 |
| GSU2609 | PilB/PulE/GspE family ATPase | 40% | 9.50 |

Description is from NCBI Reference Sequence of *G. sulfurreducens*
(https://www.ncbi.nlm.nih.gov/, GenBank accession number NC\_002939.5). Identity is based on NCBI BLAST (https://blast.ncbi.nlm.nih.gov/Blast.cgi) with PilB (GSU1491) as the query. The values (log2signal) of Transcript abundance are from those for growth on a current-producing anode (32).

### Implications

The results presented here demonstrate that deleting *pilB* is not an effective strategy for evaluating the function of e-pili in *G. sulfurreducens*. One of the primary benefits attributed to deleting *pilB* in *G. sulfurreducens* was that it yielded a strain without e-pili in which the outer-surface *c*-type cytochromes were properly localized. The same result is achieved with the expression of pilin monomers that yield poorly conductive pili (14, 20) with the added benefit that other possible functions of e-pili, such as aiding in attachment to surfaces and biofilm formation (13) are retained.

## Materials and Methods

### Strains and growth conditions

*G. sulfurreducens* strains were grown under anaerobic conditions at 30°C in a defined medium with acetate as the electron donor and fumarate as the electron acceptor as previously described (35) unless otherwise described. *Escherichia coli* NEB 10-beta (New England Biolabs) was used for plasmid preparation and grown in medium supplemented as instructed by the manufacturer, with appropriate antibiotics if necessary.

### Construction of ΔPilB strains

The *pilB* gene (GSU1491) was replaced by a kanamycin-resistance gene in *G. sulfurreducens* PCA via the double-crossover homologous recombination as described previously (35). Flanking DNA fragments were amplified by PCR with primer pairs ATC<u>TCTAGA</u>TTCCTCATAAATCGGCCATC (XbaI site is underlined)/TCT<u>GAATTC</u>AGTCTGCTAGCCTGCATAG (EcoRI site is underlined) for the upstream region and TCT<u>AAGCTT</u>CTATCAGGTAATGCCCATG (HindIII site is underlined)/TCT<u>GGTACC</u>TCGATGGTCACAATATGATC (KpnI site is underlined) for the downstream region. The *G. sulfurreducens* PCA chromosome DNA was used as the template. DNA fragment of the kanamycin-resistance gene was amplified by PCR as described previously (36). These PCR products were digested with restriction enzymes, ligated, and cloned in a plasmid. The plasmid thus constructed was linearized by XbaI. The linearized DNA fragment was used for electroporation. The deletion of the *pilB* gene and replacement with the kanamycin-resistance gene were verified by PCR with primer pairs TTCCTCATAAATCGGCCATC/GCAGTCTTGATGGACGACTC and TTCCTCATAAATCGGCCATC/ACATTCATCCCAGGTGGCAC, respectively.

The *pilB* gene (KN400_1518) was replaced by the kanamycin-resistance gene in KN400-PilA-WT/PilA-6His as described above, except that the *G. sulfurreducens* KN400 chromosome DNA was used as the template for PCR. There are 3 nucleotide differences between PCA and KN400 in the downstream region used for the recombination.

### Current production

Current production was determined in previously described bioelectrochemical systems (17) with acetate as the electron donor and positively poised (300 mV versus Ag/AgCl) graphite anodes as the electron acceptor. Once current production was initiated the anode chamber received a steady input of fresh medium as previously described (17).

### Atomic Force Microscopy

Atomic force microscopy (AFM) was carried out, with slight modification, as previously described (22). Briefly, 100 μl of a pili preparation sheared from cells was drop cast onto HOPG and allowed to set for 10 min, after which it was washed twice with 100 μl of deionized water and blotted dry. The sample was equilibrated with atmospheric humidity for at least 2 h before subsequent measurements. AFM was performed on an Oxford Instruments Cypher ES Environmental AFM in ORCA electrical mode equipped with a Pt/Ir-coated Arrow-ContPT tip with a 0.2 N/m force constant (NanoWorld AG, Neuchâtel, Switzerland). Topographical identification of the pili was achieved in contact mode, with a set point of 0.002 V. Conductive measurements were acquired using 0.002 V set point, and current-voltage curves of ±0.6 V voltage sweep at a frequency of 0.99 Hz, were generated for three spatially different points on three independent pili. Conductance was calculated by using the linear slope between −0.2 and 0.2 V.

### Transmission electron microscopy and immunogold labeling

Immunogold labeling was conducted with the 6x-His Tag polyclonal antibody as the primary antibody and the anti-rabbit IgG-gold (10 nm) antibody as the secondary antibody as previously described (26). For transmission electron microscopy 7μL of sample was dropcast on a plasma-sterilized, 400 mesh copper carbon-coated ultralight grid for 10 minutes. Excess liquid was wicked off and the grid was stained with 3μL of 2% uranyl acetate for 15-20 seconds before excess liquid was removed and air dried. Grids were examined with transmission electron microscopy on a FEI Tecnai 12 at 120kV or a JEOL 2000fx at 200kV.

## Acknowledgements

The transmission electron microscopy images were collected in the Electron Microscopy Facility at the Institute for Applied Life Sciences, UMASS-Amherst.

This research was supported by the Army Research Office and was accomplished under Grant Number W911NF-17-1-0345. The views and conclusions contained in this document are those of the authors and should not be interpreted as representing the official policies, either expressed or implied, of the Army Research Office or the U.S. Government.

## Notes

### Competing Interest Statement

The authors have declared no competing interest.

